# Incidence and Characteristics of Co-infection and Secondary Infection in Patients with COVID-19

**DOI:** 10.1101/2021.01.06.425542

**Authors:** Yingyi Guo, Jiuxin Qu, Lingling Cheng, Xiaohe Li, Ningjing Liu, Tungngai Li, Ying Jiang, Qiao Wan, Chuyue Zhuo, Shunian Xiao, Baomo Liu, Yan Chen, lin Fu, Zhixu Chen, Mingcong Ma, Chao zhuo, Nanshan Zhong

**Affiliations:** State Key Laboratory of RespiratoryDisease, First AffiliatedHospital of Guangzhou Medical University, 151 Yanjiang Rd, Guangzhou, Guangdong; Department of Infectious Diseases, The Third People’s Hospital of Shenzhen, Southern University of Science and Technology, National Clinical Research Center for Infectious Diseases, Shenzhen, China

**Keywords:** COVID-19, co-infection, secondary infection, PUATs, pathogen

## Abstract

**Objective:** The etiology and epidemiology of co-infection and secondary infection in COVID-19 patients remain unknown. The study aims to investigate the occurrence and characteristics of co-infection and secondary infection in COVID-19 patients, mainly focusing on Streptococcus pneumoniae co-infections.

**Methods:** This study was a prospective, observational cohort study of the inpatients diagnosed with COVID-19 in two designated hospitals in south China enrolled between Jan 11 and Feb 22, 2020. The urine specimen was collected on admission and applied for pneumococcal urinary antigen tests (PUATs). Demographic, clinical and microbiological data of patients were recorded simultaneously.

**Result:** A total of 146 patients with a confirm diagnosis of COVID-19 at the median age of 50.0 years (IQR 36.0-61.0) were enrolled, in which, 16 (11.0%) were classified as severe cases and 130 (89.0%) as non-severe cases. Of the enrolled patients, only 3 (2.1%) were considered to present the co-infection, in which 1 was co-infected with S.pneumoniae, 1 with B. Ovatus infection and the other one with Influenza A virus infection. Secondary infection occurred in 16 patients, with S. maltophilia as the most commonly isolated pathogen (43.8%), followed by P. aeruginosa (25.0%), E. aerogenes (25.0%), C. parapsilosis (25.0%) and A. fumigates (18.8%).

**Conclusion:** Patients with confirmed COVID-19 were rarely co-infected with Streptococcus pneumoniae or other pathogens, indicating that the application of antibiotics against CAP on admission may not be necessary in the treatment of COVID-19 cases.

## INTRODUCTION

COVID-19 has rapidly developed into a global pandemic and continues to spread rapidly all over the world.[1] According to the report released by the World Health Organization (WHO), more than 70 million people were infected with SARS-CoV-2 and meanwhile induced over 1.6 million deaths as of Dec 13 2020, bringing tremendous burden to the global society.[2]

To date, many studies have been conducted in the areas of COVID-19 epidemiology, infection characteristics, control methods and therapeutic strategies.[3] However, most microbiology laboratories are barely functioning due to the restrictions imposed on biological safety during the pandemic. Therefore, very few studies could focus on and exactly describe the problems of co-infection and secondary infection in patients with COVID-19,in which the attention on co-infection was even less.[4] Accordng to the COVID-19 treatment guidance published by WHO, early empiric antimicrobial use against co-infected community-acquired pneumonia (CAP), whose pathogen is mainly Streptococcus pneumoniae, is suggested for COVID-19 patients with severe conditions.[5] In addition, some studies suggested the application of antibacterial or antifungal medicines for coronavirus should refer to the guidelines of influenza virus infection treatment, which include early empirical antibiotics therapy.[6] This makes sense as bacterial co-infections were found in more than 40% patients with influenza and streptococcus pneumoniae was the major pathogens (>50%).[7] To date, the incidence of co-infections caused by Streptococcus pneumoniae coinciding with COVID-19 has not been adequately reported.[8] The blind empirical use of antibiotics for COVID-19 patients without pathogen detection is worrisome because it may exacerbate antimicrobial resistance (AMR).[9] Based on the few published papers, the lack of evidences of occurrence of co-infecting with streptococcus pneumoniae and other pathogens in COVID-19 patients may cause the concern for the rational use of antibiotics in the therapeutic process. [4]

This study mainly aims to investigate the incidence and characteristics of streptococcus pneumoniae co-infections in COVID-19 patients using pneumococcal urinary antigen test (PUATs), in addition to those of the secondary infection caused by other pathogens in Southern China.

## Materials and Methods

### Setting and patient

This study was designed as a prospective, observational cohort study. The study was conducted in 2 hospitals in south China, namely, the Third People’s Hospital of Shenzhen (TPHS) and Guangzhou Eighth People’s Hospital (GEPH). These are major infectious diseases hospitals in South China and were directively designated to admit COVID-19 patients by National Health Commission of the People’s Republic of China. According to Laboratory Biorisk Management for Laboratories Handling Human Specimens Suspected or Confirmed to Contain Novel Coronavirus 2012 Interim Recommendations published by WHO, routine examination of fungal and bacterial cultures developed from respiratory tract specimens were performed in BSL2 laboratory with BSL3 standard precautions. Both hospitals met the standard so that routine pathogen detection could be performed.[10]

### We enrolled the patients with confirmed COVID-19 admitted between Jan 11 to Feb 22, 2020. Microbiology

Patients underwent microbiological examinations of bacteria, fungi and common respiratory viruses within 48 hours of admission. Etiological detection of bacteria and fungi were conducted though PUATs(BinaxNOW) and cultures. After obtaining the informed consents of all enrolled patients, urine samples were collected on admission to accomplish PUATs. The culture tests were performed with blood, urine and sputum. Blood culture was performed on patients with body temperature exceeding 37.3°Cand sputum culture was performed when symptoms of expectoration were exhibited. If there were signs related to urinary system infection, urine culture would be performed. The detection tests of common respiratory viruses such as Legionella pneumophila, Mycoplasma pneumoniae, Q fever Rickettsia, Chlamydia pneumoniae, adenovirus, respiratory syncytial virus, influenza A virus, influenza B virus and parainfluenza virus types 1, 2 and 3 were also performed in all patients by Polymerase Chain Reaction (PCR).

The microbiological examinations done after 48 hours of admission were mainly cultures, including blood, urine, sputum and bronchoalveolar lavage fluid (balf) culture, that were conducted in accordance with Diagnosis and Treatment of Community-acquired Pneumonia in Adults: 2016 Clinical Practice Guidelines released by the Chinese Thoracic Society, Chinese Medical Association.[10]

### Data collection and assessment

Patients’ information from admission to discharge was collected from electronic medical records. Demographic characteristics including sex, age and comorbidities were recorded. Clinical manifestations were collected including symptoms, laboratory findings, antibiotics exposure, length of Intensive Care Unit (ICU) or hospital stay, mechanical ventilation administration as well as the prognostic status (recovery or death), and microbiological test results including type of specimen, time of pathogen isolation and pathogens detected. All data were reviewed, recorded and cross-checked by 3 experienced respiratory professionals. If one of them had different opinion, opinions of the other two would be still adopted. If three of them had disagreements, an independent senior clinician woud be invited for evaluation.

### Diagnosis of COVID-19, co-infection and secondary infections

Patients who tested positive for high-throughput sequencing or real-time reverse transcription polymerase-chain-reaction on nasal or pharyngeal swab samples were diagnosed as COVID-19 patients. The COVID-19 patients are classified as severe cases or non-severe cases based on the American Thoracic Society guidelines for community-acquired pneumonia, which has been cited in many previous studies.[8] The diagnosis of severe case should meet one major criterion or more than three minor criteria.[11] Considering the fact that COVID-19 is community-acquired pneumonia, community-acquired pathogens in addition to SARS-COV-2 are considered co-infection, while hospital-acquired pathogens are considered secondary infections. Co-infection cases are patients with positive PUATs, with at least one positive results of blood, sputum and urine cultures or one common respiratory virus within 48 hours of admission.[12] Secondary infections are patients with positive culture of new pathogens from blood, sputum, balf and urine samples 48 hours after admission.[4] Bacteremia was determined in patients with positive blood cultures.[13]

### Statistical analysis

All statistical analysis was completed using SPSS v.25.0. Continuous variables were reported as means and standard deviations (SD) for normally distributed data or as medians and interquartile range (IQR) for abnormally distributed data. Categorical variables were reported as frequencies and percentages. Kolmogorov-Smirnov test was used to identify the normality of data. Comparisons of categorical variables between groups were made using Person Chi-Square or Fisher’s Exact Test. Comparisons of continuous variables were performed by Student’s T-test when normality was demonstrated; otherwise, Mann-Whitney U was used. All tests were two-tailed, and the statistical significance was set at P < 5%.

## RESULTS

### Demographic, clinical, laboratory findings and cilinical outcomes

The data of baseline demographic characteristics, clinical manifestations and laboratory findings were summarized in Table 1. During the study period, 146 patients with a confirmed diagnosis of COVID-19 were admitted into the two hospitals. The median age was 50.0 years (IQR 36.0-61.0) and among them 69 patients (48.6%) were male. Hypertension (28, 19.2%) was the most common comorbidity, followed by diabetes (11, 7.5%) and coronary heart disease (8, 5.5%). Cough (76, 52.1%) and fever (41, 28.1%) were the most seen clinical manifestations. The median length of hospital stay was 19 (IQR 13-26) days. 27 patients were transferred to ICU and the median length of ICU stay was 15.5 (IQR 9.5-28.5) days. Based on the American Thoracic Society guidelines for community-acquired pneumonia, 16 (11.0%) patients were considered as severe cases and 130 (89.0%) as non-severe cases. The white blood cell(WBC) count of 39 (26.7%) patients was below the normal range and that of 7 (4.8%) patients was above, Lactate dehydrogenase(LDH) and D-dimer,were abnormal in 65 (44.5%) patients, and 96 (65.8%) patients, respectively. Only one patient showed a higher procalcitonin(PCT) level than the normal range. The median level of interleukin-6 (IL-6) and erythrocyte sedimentation rate (ESR) are 10.99 (3.72-24.98) and 30.0 (14.5-55.0). Imaging abnormalities were shown in 140 (95.9%) patients. Three (2.1%) patients were dead eventually.

**Table 1.**
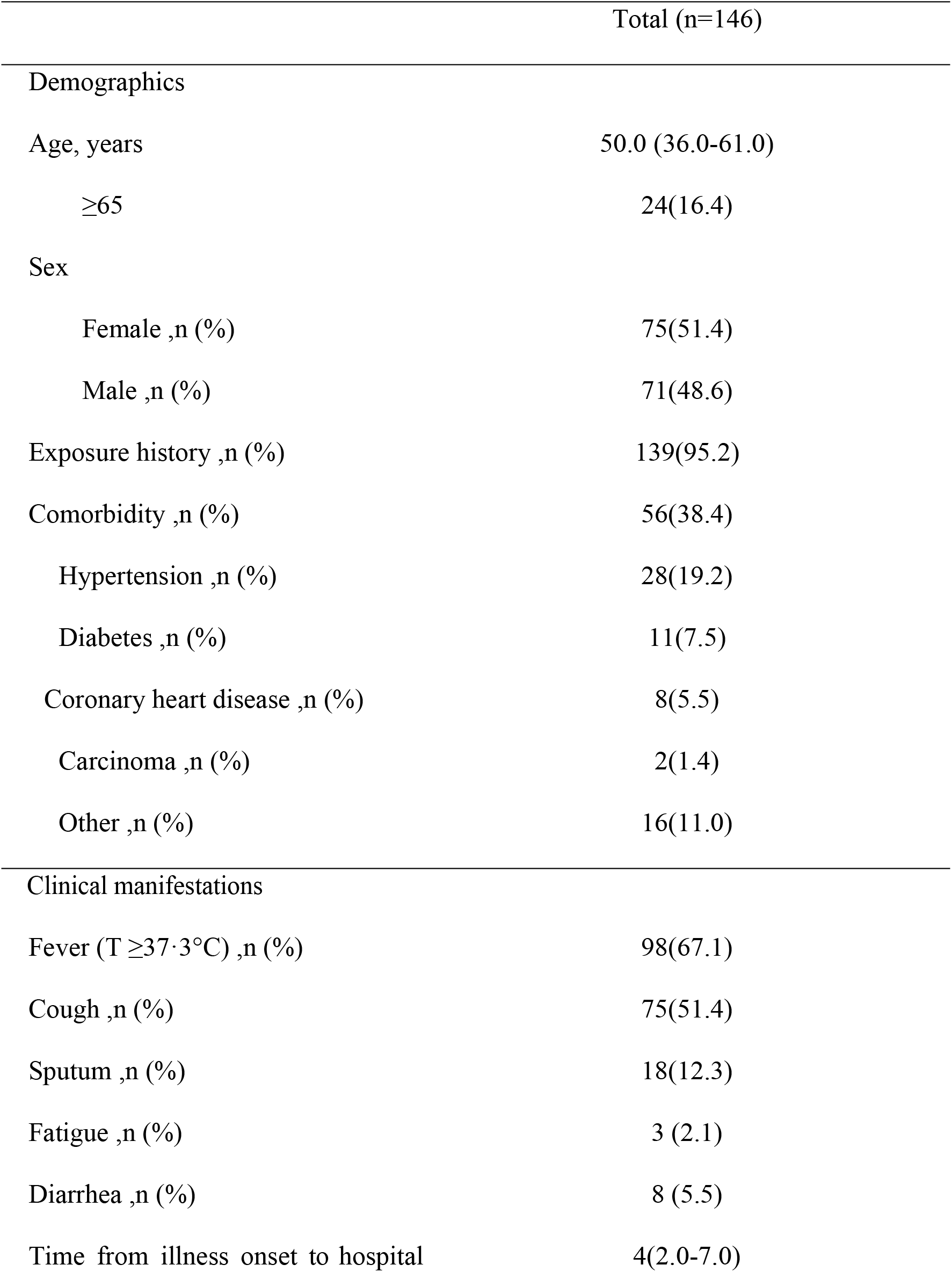
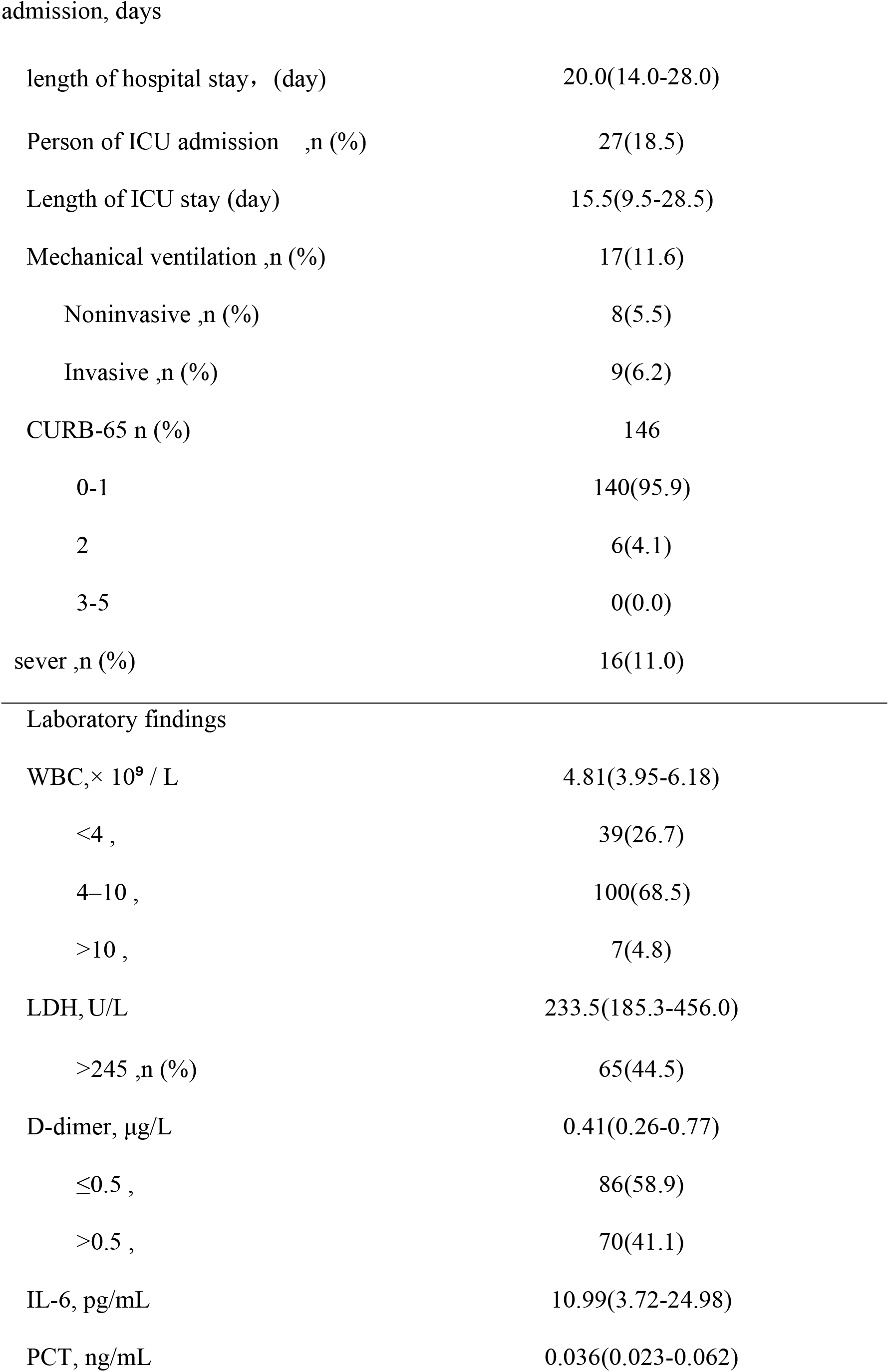
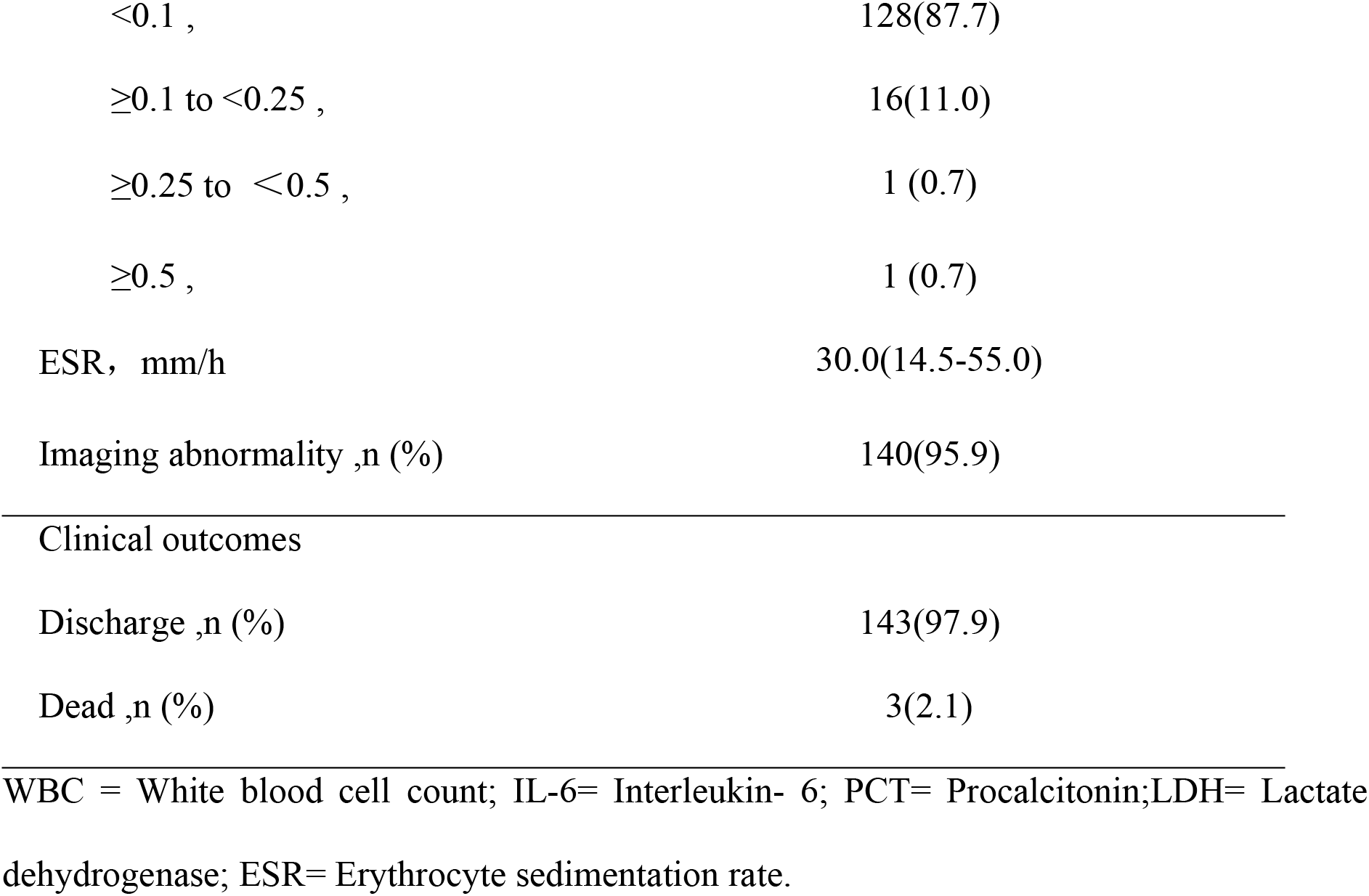
Baseline Demographic Characteristics, Clinical Manifestations, Laboratory Findings and Clinical Outcomes in Severe and non-Severe COVID-19 patients [Median (IQR) or n]

|  | Total (n=146) |
| --- | --- |
| Demographics |  |
| Age, years | 50.0 (36.0-61.0) |
| $\geq 65$ | 24(16.4) |
| Sex |  |
| Female ,n (%) | 75(51.4) |
| Male ,n (%) | 71(48.6) |
| Exposure history ,n (%) | 139(95.2) |
| Comorbidity ,n (%) | 56(38.4) |
| Hypertension ,n (%) | 28(19.2) |
| Diabetes ,n (%) | 11(7.5) |
| Coronary heart disease ,n (%) | 8(5.5) |
| Carcinoma ,n (%) | 2(1.4) |
| Other ,n (%) | 16(11.0) |
| Clinical manifestations |  |
| Fever ($T \geq 37.3^{\circ}\text{C}$) ,n (%) | 98(67.1) |
| Cough ,n (%) | 75(51.4) |
| Sputum ,n (%) | 18(12.3) |
| Fatigue ,n (%) | 3 (2.1) |
| Diarrhea ,n (%) | 8 (5.5) |
| Time from illness onset to hospital | 4(2.0-7.0) |
| length of hospital stay, (day) | 20.0(14.0-28.0) |
| Person of ICU admission ,n (%) | 27(18.5) |
| Length of ICU stay (day) | 15.5(9.5-28.5) |
| Mechanical ventilation ,n (%) | 17(11.6) |
| Noninvasive ,n (%) | 8(5.5) |
| Invasive ,n (%) | 9(6.2) |
| CURB-65 n (%) | 146 |
| 0-1 | 140(95.9) |
| 2 | 6(4.1) |
| 3-5 | 0(0.0) |
| sever ,n (%) | 16(11.0) |
| Laboratory findings |  |
| WBC,× 10<sup>9</sup> / L | 4.81(3.95-6.18) |
| <4 , | 39(26.7) |
| 4–10 , | 100(68.5) |
| >10 , | 7(4.8) |
| LDH,U/L | 233.5(185.3-456.0) |
| >245 ,n (%) | 65(44.5) |
| D-dimer, µg/L | 0.41(0.26-0.77) |
| ≤0.5 , | 86(58.9) |
| >0.5 , | 70(41.1) |
| IL-6, pg/mL | 10.99(3.72-24.98) |
| PCT, ng/mL | 0.036(0.023-0.062) |
| <0.1 , | 128(87.7) |
| ≥0.1 to <0.25 , | 16(11.0) |
| ≥0.25 to <0.5 , | 1 (0.7) |
| ≥0.5 , | 1 (0.7) |
| ESR, mm/h | 30.0(14.5-55.0) |
| Imaging abnormality ,n (%) | 140(95.9) |
| Clinical outcomes |  |
| Discharge ,n (%) | 143(97.9) |
| Dead ,n (%) | 3(2.1) |

WBC = White blood cell count; IL-6= Interleukin- 6; PCT= Procalcitonin;LDH= Lactate dehydrogenase; ESR= Erythrocyte sedimentation rate.

### Microbiology tests and characteristics of patients with co-infection

The pathogens detected in co-infection patients were summarized in Table 2. Of 146 patients, 3 (2.1%) were considered to present co-infection, in which 1 was co-infected with S.pneumoniae confirmed, 1 with B. Ovatus and the other one with Influenza A virus infection. PUATs was conducted among all 146 patients with 1 positive case (0.7%). Blood, urine and sputum culture were performed in 2, 1 and 48 patients respectively within 48 hours of admission; among them, only one patient had a positive bacterial culture result (B. Ovatus positive in blood culture). Of the 146 patients experienced common respiratory viruses tests on admission, only one (0.7%) patient was positive for influenza A virus test.

**Table 2.**
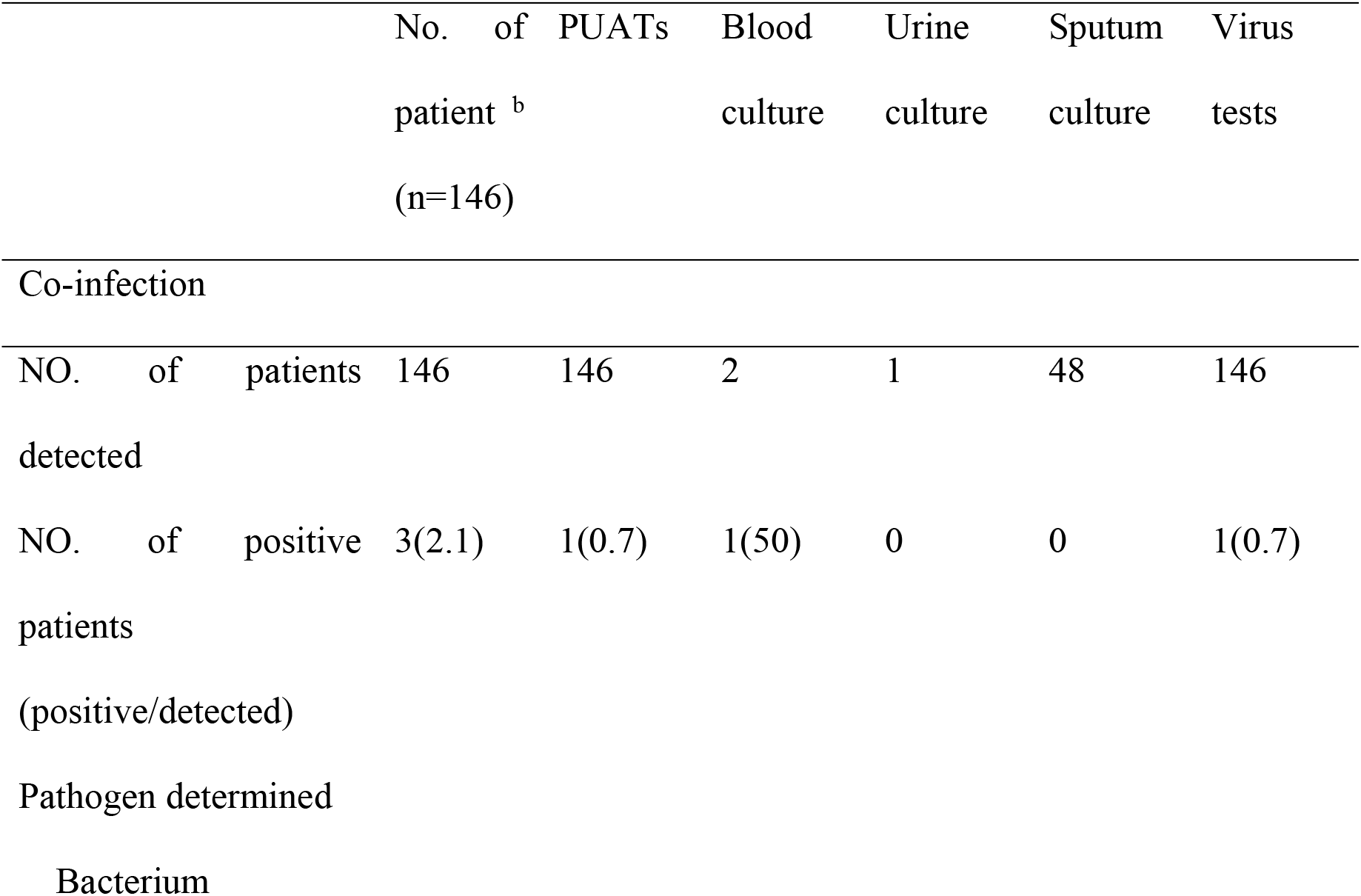
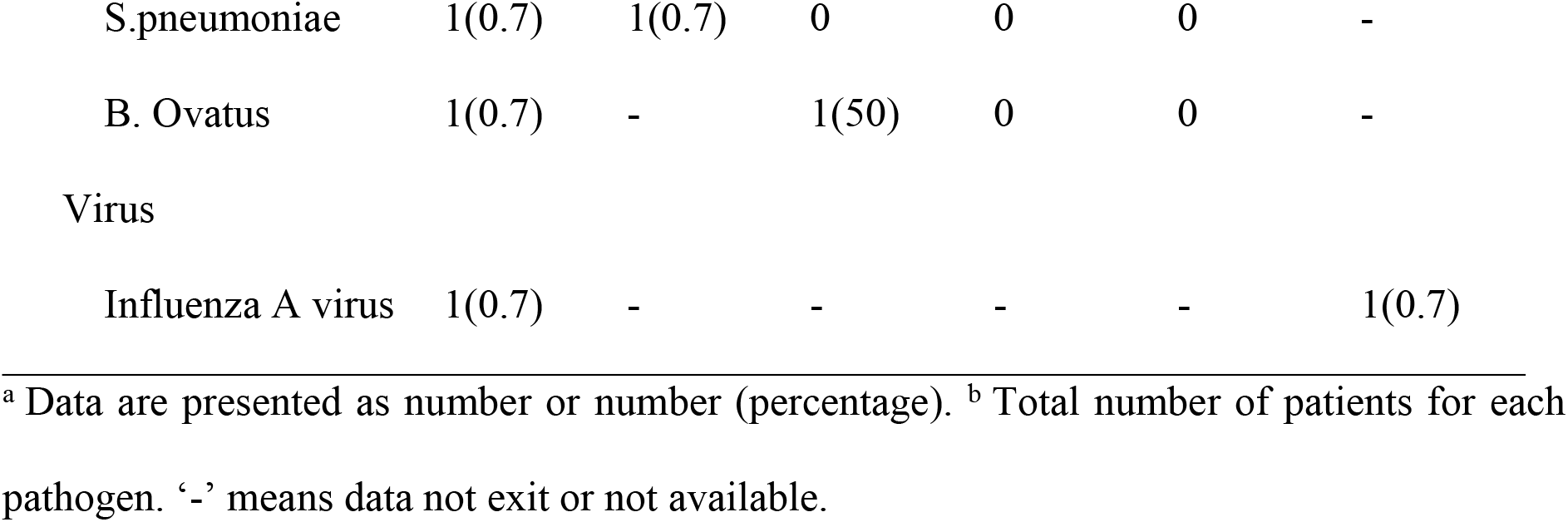
Microbiology tests of co-infection ^a^[Median (IQR) or n]

Clinical Information of three patients with co-infection was listed in Table 3. All three patients with co-infection were male and the age was 72, 49 and 63 years old, respectively. In the 3 patients with co-infection, 2 patients (1 with S. pneumoniae and 1 with Influenza A Virus) were considered as non-severe cases and 1 patient were considered as severe case, who died from B. Ovatus co-infection. Because of the rare cases of co-infection, univariate analysis was not performed.

**Table 3.** clinical Information about patients with co-infection

|  | Patient 1 | Patient 2 | Patient 3 |
| --- | --- | --- | --- |
| Co-infected pathogen | S. pneumoniae | Influenza A Virus | B. Ovatus |
| Sex | male | male | male |
| Age, years | 72 | 49 | 63 |
| Comorbidity | Hypertension | other | other |
| Clinical symptoms | Fever,cough | Fever | Fever, cough,<br>thoracodynias,<br>polypnea |
| White blood cell<br>count,<br>× 10<sup>9</sup> per L | 8.42 | 3.77 | 6.79 |
| IL-6, pg/mL | 15.65 | 79.45 | 3.17 |
| Lactate<br>dehydrogenase, U/L | 200 | 139 | 720 |
| D-dimer, µg/L | 2.30 | 0.28 | 2.85 |
| Procalcitonin, ng/mL | 0.03 | 0.04 | 9.18 |
| ESR, mm/h | 3 | 14 | 2 |
| Imaging features | Abnormal | Abnormal | Abnormal |
| Mechanical Ventilation | - | - | Invasive mechanical ventilation |
| Disease severity status | Common | Common | Critic |
| Secondary infection | No | No | Yes |
| Admission to ICU, yes or no | No | No | Yes |
| Use of antibiotics | Yes | No | Yes |
| Outcome | discharge | discharge | dead |

### Microbiology tests and analysis of patients with secondary infection

The pathogens detected in secondary infection patients were summarized in Table 4.

**Table 4.**
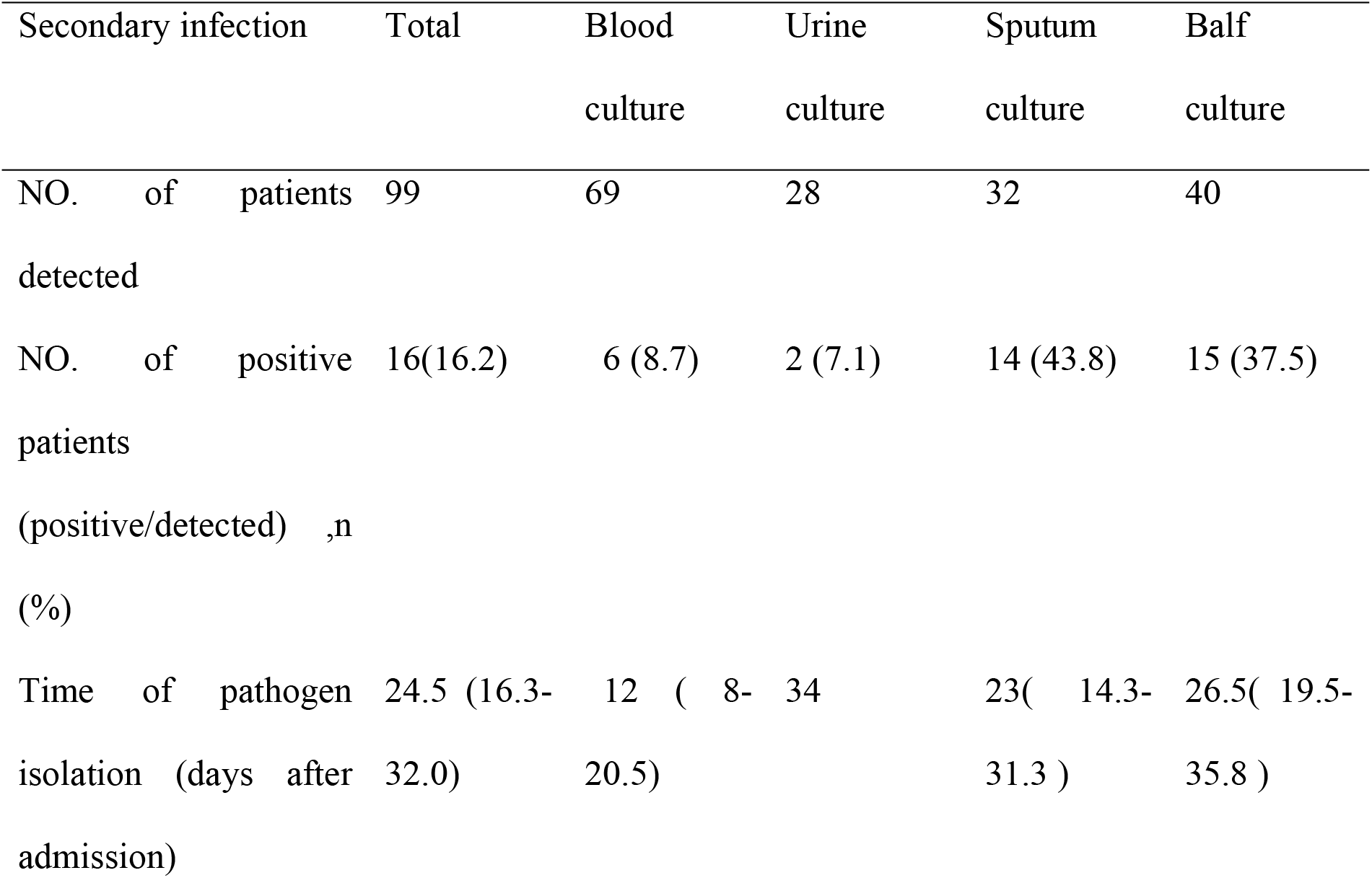
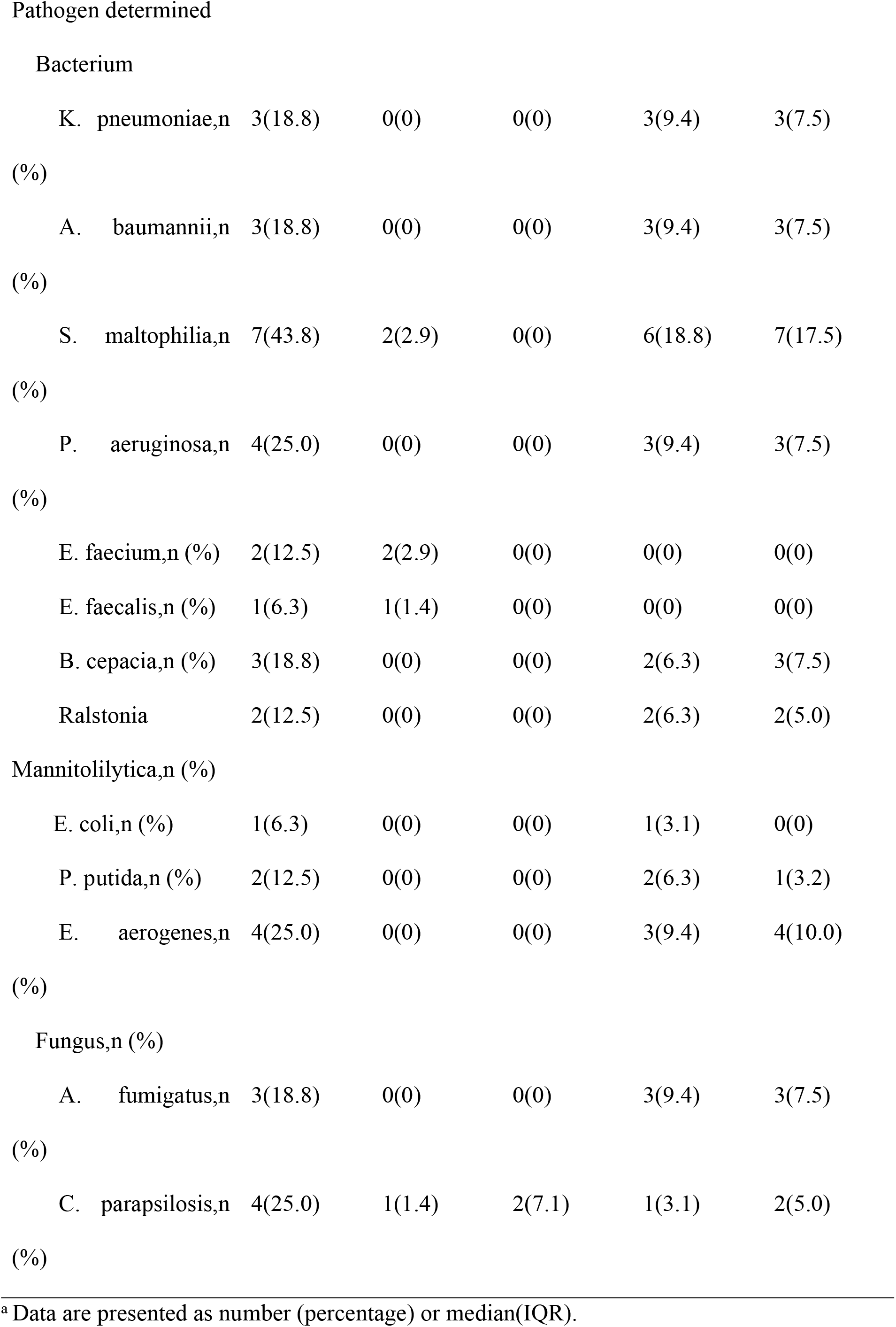
Microbiology tests of secondary infection ^a^[Median (IQR) or n]

In this study, 99 patients underwent at least one of the following tests including blood culture, urine culture, sputum culture and bronchoalveolar lavage fluid culture 48 hours after admission. Among them, 16 (16.2%) patients showed positive results for at least one pathogen besides SARS-COV-2 and were regarded as secondary infection. Blood, urine, sputum and balf culture were performed in 69, 28, 40 and 32 patients with the positive results in 6 (8.7%), 2 (7.1%), 15 (37.5%) and 14 (43.8%) patients, respectively. The median time of pathogens isolated was 24.5 (16.3-32.0) days after admission.

Of the 16 patients with secondary infection, 15 (93.8%) showed positive bacterial culture and 6 (37.5%) showed positive fungal culture.

In secondary bacterial infections, S. maltophilia was the most common pathogen (43.8%), followed by P. aeruginosa (25.0%), and E. aerogenes (25.0%). K. pneumoniae, A. baumannii and E. coli were always blamed in nosocomial infection and accounted for 18.8%, 18.8%, 6.3%, respectively. Staphylococcus aureus were not detected. Five (31.3%) patients with secondary infection showed positive bacteria blood culture and were considered as bacteremia (S. maltophilia in 2 cases, E. faecium in 2 cases, and E. faecalis in 1 case).

Of the 16 patients with secondary infection, 6 cases (37.5%) showed positive results in fungal culture (3 for A. fumigatus and 4 for C. parapsilosis, and 1 for both); in which, 1 case showed positive blood culture of C. parapsilosis, which was considered as fungemia.

The data of patients with or without secondary infection were demonstrated in Table 5. Patients with secondary infection were usually older, co-existing more and severe diseases, and more likely to have diarrhea than those without secondary infection (P < 0.05). Also, there are more patients admitted to ICU and mechanically ventilated as well as longer length of hospital and ICU stay among patients with secondary infection (Table 5, P < 0.05). As to disease severity, patients with secondary infection appeared to be more severe.

**Table 5.**
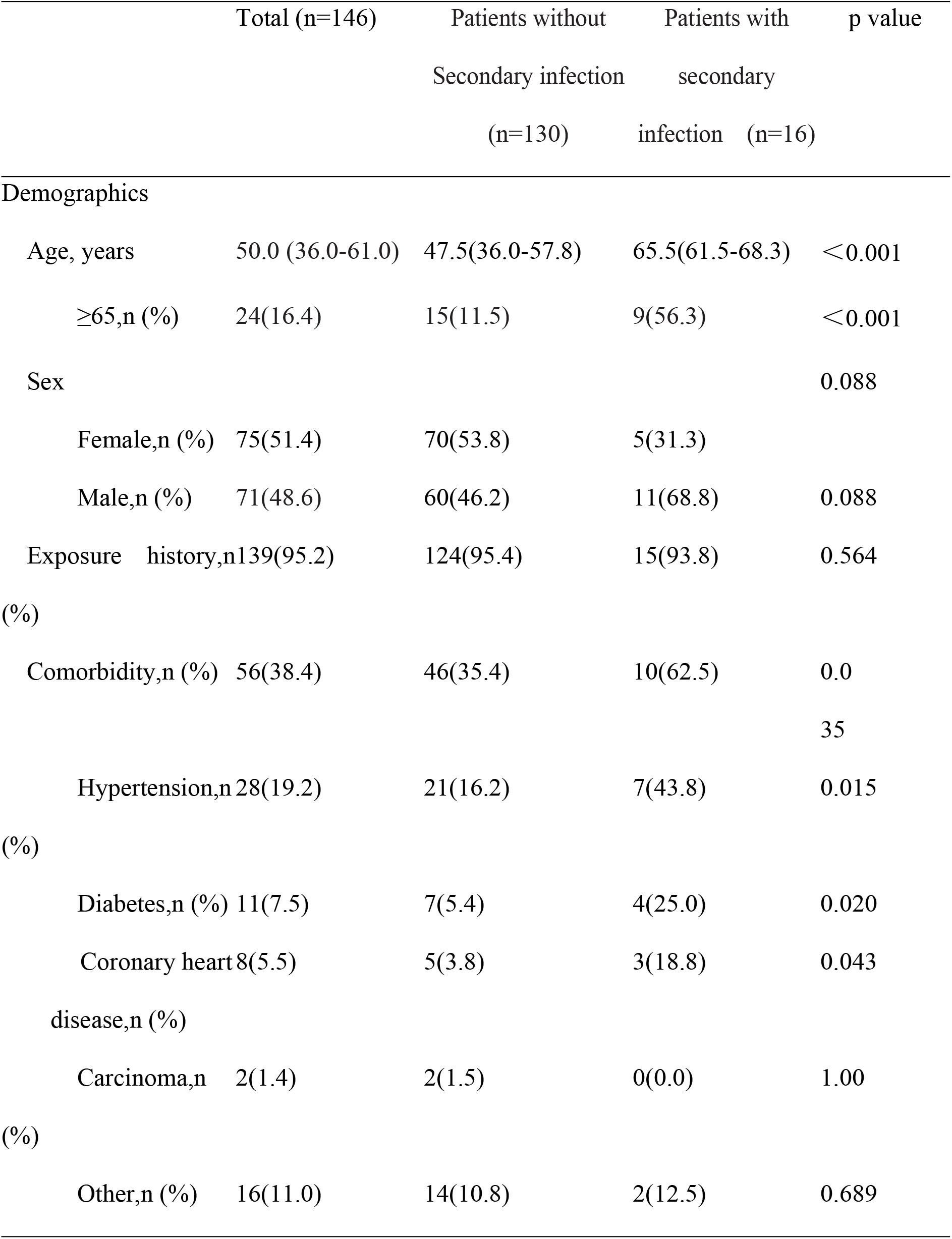
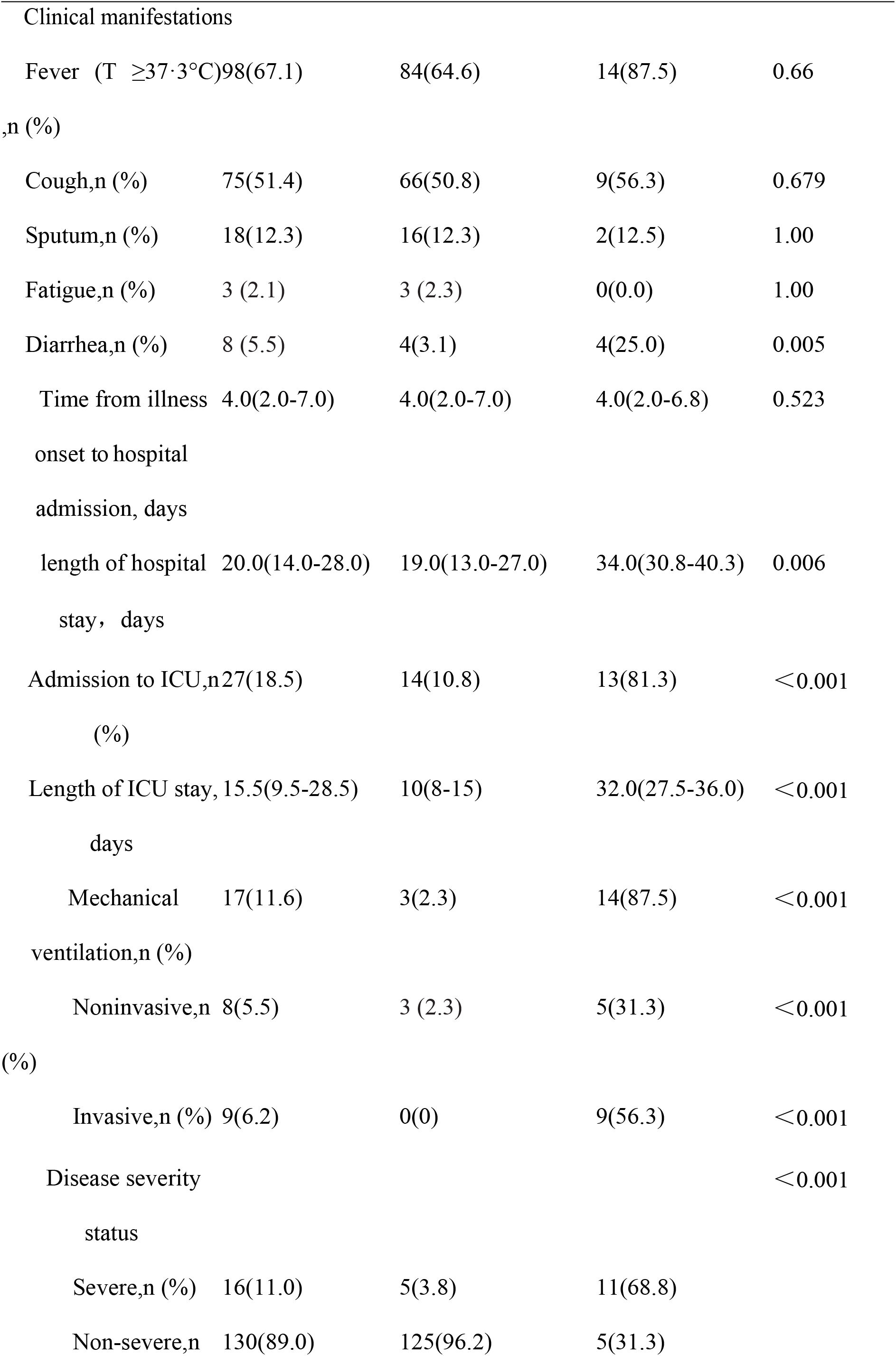
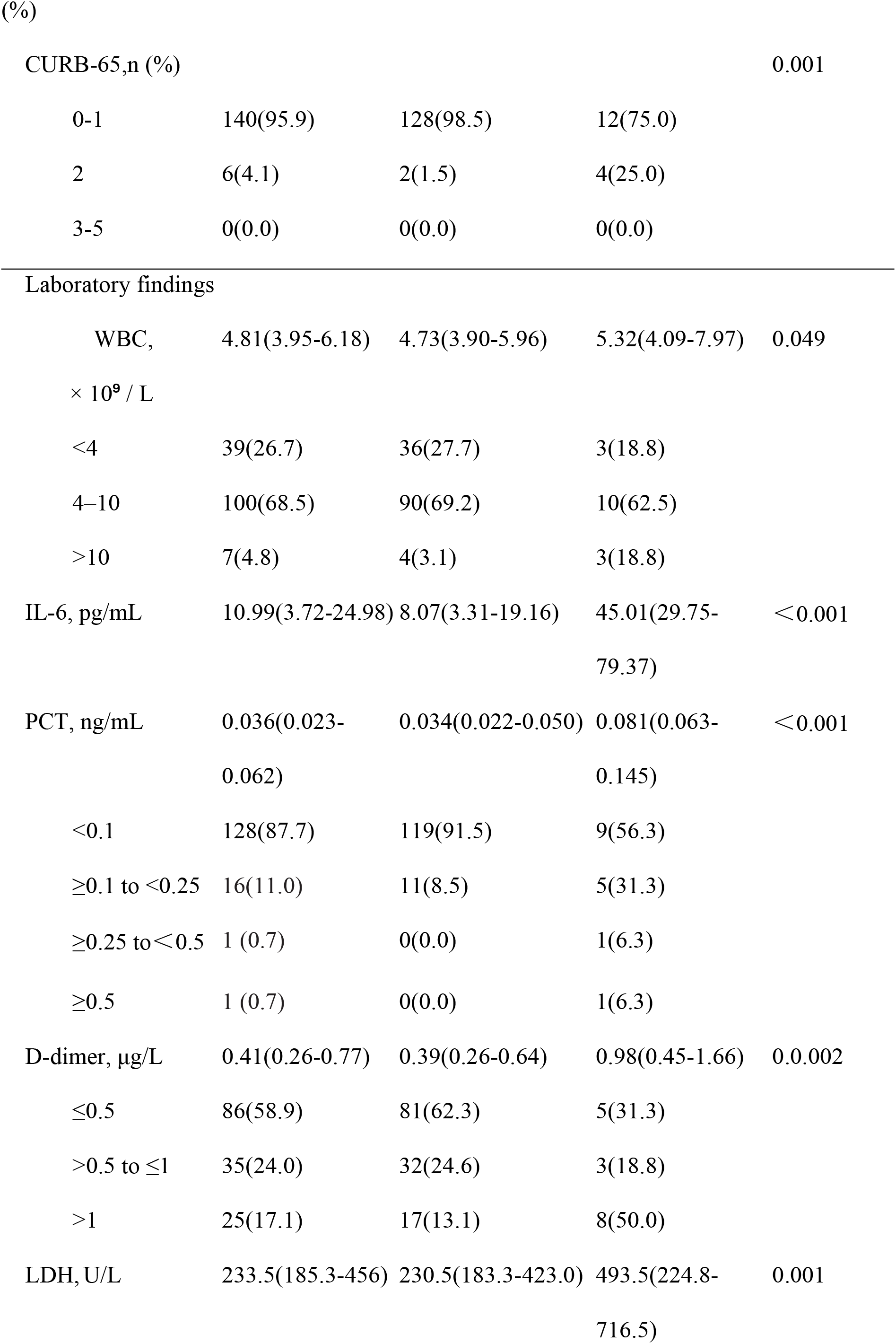
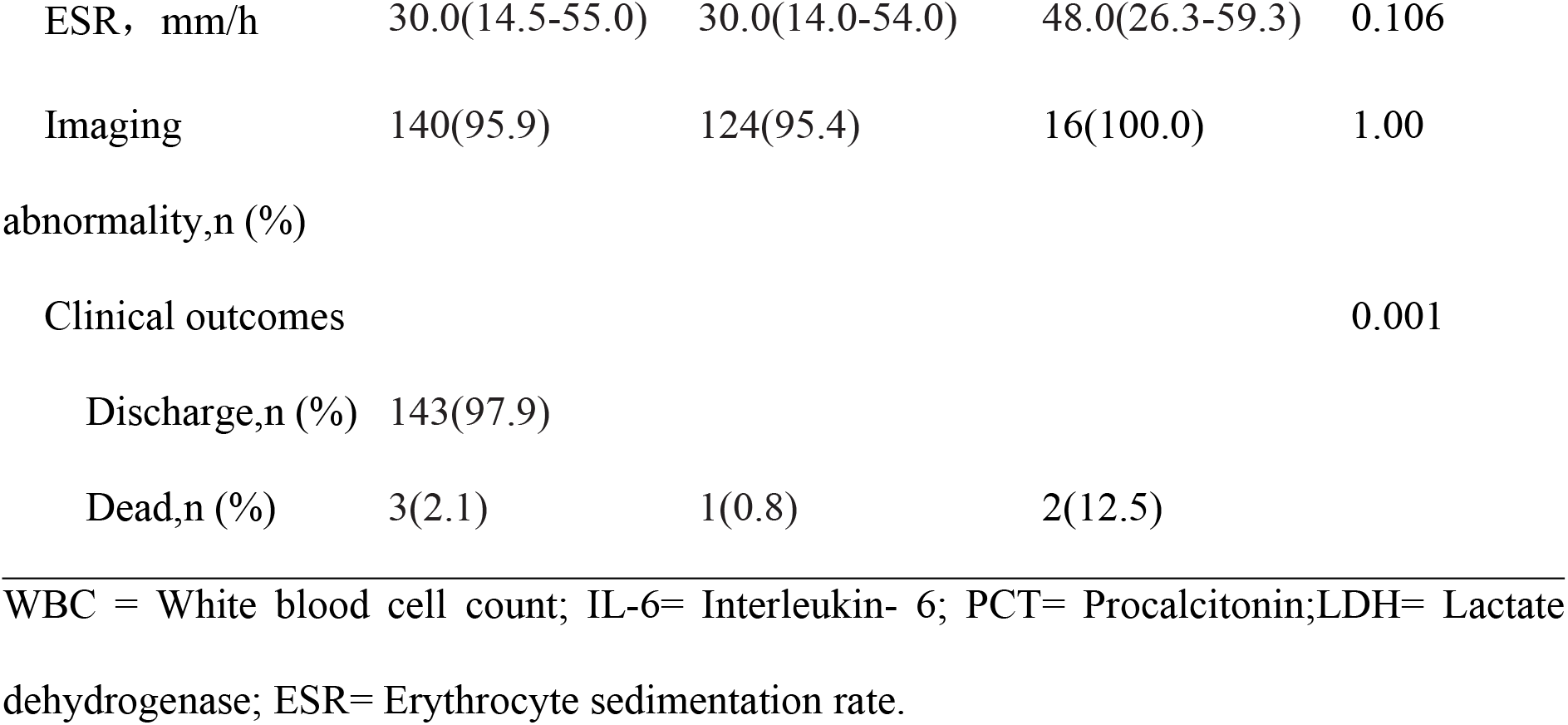
Comparison of demographic characteristics, clinical manifestations and laboratory findings between COVID-19 patients with or without secondary infection[Median (IQR) or n]

Major laboratory markers were tracked from the time of admission. For patients with secondary infection, baseline IL-6, procalcitonin, CRP, ESR were significantly higher(table 5). In this study, all three patients who eventually died were accompanied by positive results of blood culture (2 with secondary S. maltophilia bacteremia and 1 with co-infected B. ovatus bacteremia), suggesting the mortality of patients with bacteremia reaching 42.9% (3/7).

## DISCUSSION

As the COVID-19 pandemic continues, whether COVID-19 patients need regular empirical antibiotic treatment has become a big concern. Some studies suggested that the influenza treatment guidelines can be referred for the treatment of bacterial infections in COVID-19 patients.[6] However, the lack of recognition of co-infections or secondary infections in COVID-19 patients impedes us from seeing whether their patterns are similar to those of flu. This study disclosed the pattern of co-infections induced by Streptococcus pneumoniae and other pathogens in patients with COVID-19. Contrary to the high occurence rate of Streptococcus pneumoniae co-infections with influenza, our results showed that there is a extremely low incidence of co-infection caused by Streptococcus pneumoniae among patients with COVID-19 (0.7%). Besides, co-infections caused by other pathogens were rare (1.4%) too.

In our study, only 2 bacterial and 1 viral co-infection cases were found. No fungal co-infection was detected. PUAT was used for Streptococcus pneumoniae co-infections detection on all subjects in this study; but only 1 (0.7%) case showed positive result. Given the fact that the sensitivity and specificity of PUAT range from 59-87% and 80-100% respectively, it is possible that few positive results may be missing.[14] Besides, Stenotrophomonas maltophilia was detected in only 1 patient’s blood culture sample while his other culture tests were negative. Previous research reported that PCT level could indicate the presence of bacterial co-infections in viral pneumonia.[15] Apart from the patients with bacteremia co-infected mentioned above, PCT levels of all patients were within the normal range. Considering the results of PUAT, culture tests and PCT levels, we reckon that co-infections in COVID-19 patients were seldom seen, given the fact that few studies involving COVID-19 co-infections caused by Streptococcus pneumoniae or other pathogens were published.[4, 8, 10] Besides, studys associated with pneumonia caused by other coronavirus, such as Severe Acute Respiratory Syndrome (SARS) and Middle East Respiratory Syndrome Coronavirus (MERS), also have low occurance rate of co-infection, further supporting our conclusion.[4, 16-17] In addition to bacterial co-infection, one patient was tested positive for influenza A virus. Co-infection of influenza A with COVID-19 is occasionally reported, which is consistent with the low infection rate in this study.[18-20] Among the 3 cases described above, the patient with positive blood culture was dead and the other two with minor or moderate conditions were discharged finally.

For secondary infection, we revealed that it occurred in 16.2% COVID-19 patients, similar to the incidence rate reported in previous study.[8] After dividing our patients into subgroups based on whether they have secondary infections, we observed that patients with senior age and multiple complications were more susceptible to infection due to their weakening immune capability. Our results also disclosed that patients with secondary infections usually had severely illness and required more mechanical ventilations and longer ICU and hospital stay. The degree of disease severity was critical in COVID-19 patients with secondary infections, consistent with the observations reported in previous articles.[4] In a recent analysis of 191 cases of Wuhan Jinyintan Hospital, there was only 1 case with secondary infection in the survival group (1%, 1/131), while 50% (27/54) in the death group with unknown pathogen.[4] Recent studies have shown that the severity of patients’ disease could be revealed by molecular baseline indicators such as D-dimer and LDH, [11]which is consistent with the results of patients with secondary infection in this study.

It should be noted that bacteremia may cause a high mortality in COVID-19 patients (42.9%). All 3 patients with bacteremia died, two of whom were induced by S. maltophilia. We considered this uncommon pathogen is not representative because of its very finite prevalence in the environment and the patients’ poor immunity.

We inferred that the patterns of co-infection and secondary infection in COVID-19 patients were different from influenza viruses. COVID-19 patients rarely co-infected with S. pneumoniae and other pathogens, while influenza viruses showed a relatively high proportion of bacterial infections or other respiratory viruses infections, and the co-infection pathogens mainly included S. pneumoniae (> 50%), S. aureus, and H. influenza.[7] Our wider analysis of the epidemiological data of SARS (2003), H1N1 (2009), MERS (2014) and China Influenza (2018) suggested that influenza viruses and coronaviruses (SARS, MERS and COVID-19) had distinctive patterns of bacteria coinfection. There were few related reports of co-infection of coronaviruses. Among the limited reports, it is rare to see S. pneumoniae, S. aureus, and H. influenzae as seen in Influenza viruses.[16] As for the patterns of secondary infections, reports of nosocomial infections of coronaviruses were significantly less than those of influenza viruses. [4, 17]

It is unknown whether different pathogenesis can explain the difference between influenza virus and coronavirus co-infection and secondary infection. A recent report indicated that unlike influenza, viral sepsis played a crucial role in COVID-19 with less possibility of co-infections and/or secondary infections.[21] Therefore, it is inappropriate to apply antibacterial or antifungal medicine for coronavirus based on the guidelines for influenza virus infection treatment.

There are significant advantages in our study. Several problems on the setting and assays causing the current lack of co-infection data in COVID-19 patients were resolved in our research. During the outbreak of COVID-19, considering the biological safety of the specimen, routine bacteria smears and cultures were restricted in most microbiology laboratories, inducing difficulty in detecting co-infection with bacteria and fungi in COVID-19. The microbiological laboratories in the two hospitals designated in our study met the conditions for continuing routine microbiological testing, making it possible to understand the pathology and incidence of co-infection and secondary infection with COVID-19. On the other hand, the application of PUATs improved the chances of acquiring reliable positive results on patients co-infected with S. pneumoniae. As the early symptoms of COVID-19 patients are mostly cough without sputum, it is hard to obtain respiratory specimens, and throat swab culture cannot be used as evidence of lower respiratory infection. PUATs could circumvent this restriction by using urine samples. In addition, while the empirical use of broad-spectrum antibiotics might cause false negative results in culture tests of S. pneumoniae, the use of antibacterial drugs has limited impact on PUATs, as is evidenced by the 42% positive rates of PUATs among CAP patients who had received antibiotics prior to sample collection.[22] Furthermore, compared with using PCR method for diagnosis, PUATs target at the presence of S. pneumoniae by avoiding detection of colonized bacteria. Last but not least, PUAT could be performed on a bench under proper precautions and without biological safety cabinets since it meets the criteria of POC (Point of care or near-POC), making the screening process of S. pneumoniae co-infection much more convenient.[10]

Limitations existed also in our study. First, the empirical use of broad-spectrum antibiotics at the time of admission might produce the difficulty in detection of sensitive bacterial strains and induce the false negative results. Nevertheless, the effects of antibiotics could be neglected in this study since only 20.7% patients with negative etiology test had used antimicrobial drugs, indicating a restrained use of broad-spectrum antibiotics. Also, the low PCT levels in most patients suggest the negative results in our study are credible. Second, since this study was conducted in south China, the generalizability may be limited. Further study should be condcuted in more regions. Third, the sample size was too small to do more statistical analysis such as multivariate analysis to discover more eloquent difference among patients with or without co-infection and secondary infection. Further studies with more patients are required.

## Conclusion

Our study suggested that patients with confirmed COVID-19 are rarely co-infected with streptococcus pneumoniae or other pathogens, This indicates that the application of antimicrobials against CAP on admission may not be necessary. Patients with COVID-19 need to be regularly assessed to aviod secondary infections and receive timely diagnosis and treatment.

## Contributions

CZ and NZ designed this study. XL and YL collected data and participated in data interpretation. YG, TL and NL analyzed the data and developed a draft manuscript, and all authors participated in writing subsequent drafts. MH and KMS made the figures. All authors approved the final version of the manuscript.

## Acknowledgement

We thank Guanyang Zou for his kind advice to the manuscript. We thank all the patients in this study. We thank all the front-line medical staff in the Department of Infectious Diseases of the Third People’s Hospital of Shenzhen.

## Availability of data and materials

The datasets used and/or analysed during the current study are available from the corresponding author on reasonable request

## Ethics approval and consent to participate

This study was approved by both the Ethics Committees of the First Affiliated Hospital of Guangzhou Medical University and Shenzhen Third People’s hospital (Document Document 2020-49th).

## Consent for publication

Not applicable.

